# The kinetochore protein Spc105, a novel interaction partner of LaeA, regulates development and secondary metabolism in *Aspergillus flavus*

**DOI:** 10.1101/487280

**Authors:** Qing-Qing Zhi, Lei He, Jie-Ying Li, Jing Li, Zhen-Long Wang, Guang-Yao He, Zhu-Mei He

## Abstract

Nuclear protein LaeA is known as the global regulator of secondary metabolism in *Aspergillus*. LaeA connects with VeA and VelB to form a heterotrimeric complex, which coordinates fungal development and secondary metabolism. Here, we describe a new interaction partner of LaeA, the kinetochore protein Spc105, from the aflatoxin-producing fungus *Aspergillus flavus*. We showed that in addition to involvement in nuclear division, Spc105 is required for normal conidiophore development and sclerotia production of *A. flavus*. Moreover, Spc105 positively regulates the production of secondary metabolites such as aflatoxin and kojic acid, and negatively regulates the production of cyclopiazonic acid. Transcriptome analysis of the *Δspc105* strain revealed that 23 backbone genes were differentially expressed, corresponding to 19 of the predicted 56 secondary metabolite gene clusters, suggesting a broad regulatory role of Spc105 in secondary metabolism. Notably, the reduced expression of *laeA* in our transcriptome data led to the discovery of the correlation between Spc105 and LaeA, and double mutant analysis indicated a functional interdependence between Spc105 and LaeA. Further, yeast two-hybrid (Y2H) and GST pull-down assays revealed that Spc105 interacts directly with the *S*-adenosylmethionine (SAM)-binding domain of LaeA, and that the leucine zipper motif in Spc105 is required for this interaction. The Spc105-LaeA interaction identified in our study indicates a cooperative interplay of distinct regulators in *A. flavus*, providing new insights into fungal secondary metabolism regulation networks.

**Author summary:** The worldwide concern on agriculture and food safety due to aflatoxin contamination has prompted increasing interest in understanding aflatoxin biosynthesis regulation that has become a model for exploring secondary metabolism regulation in fungi. In *Aspergillus*, the nuclear protein LaeA is considered as a global regulator of secondary metabolism that forms a regulatory unit with VeA and VelB. LaeA is proposed to control secondary metabolism epigenetically through altering the chromatin structure for gene expression, however, the exact mechanism is still enigmatic. Here we identified a new interaction partner of LaeA, a kinetochore protein Spc105, in aflatoxigenic *Aspergillus flavus*. We presented a regulatory role of this kinetochore protein in fungal development and aflatoxin biosynthesis for the first time, and demonstrated its interdependent relationship with LaeA. The finding that such a conserved cell division related gene regulates *A. flavus* secondary metabolism will offer new insights into the fungal secondary metabolism regulation and improve our understanding of LaeA involved networks.

## Introduction

*Aspergillus flavus* is a ubiquitous saprophytic filamentous fungus that infects economically important crops such as peanuts, maize, and many other seed crops during preharvest or storage [1]. Infestations of *A. flavus* in crops are spread by the production and dissemination of airborne conidia or resistant structures called sclerotia. *A. flavus* can produce several types of mycotoxins known as aflatoxins (AF), aflatrem, and cyclopiazonic acid (CPA). AF is the most toxic and carcinogenic secondary metabolite, with its contamination resulting in huge economic losses and threatening human health [2,3]. Therefore, studies on AF biosynthesis regulatory mechanisms is vital for the development of strategies to control mycotoxin contamination.

The *A. flavus* genome has been predicted to contain 56 secondary metabolite clusters, with each cluster containing ‘backbone’ enzyme gene(s) essential to secondary metabolite biosynthesis [4,5]. The AF cluster, containing approximately 30 genes and two pathway-specific regulatory genes (*aflR* and *aflS*), has been well characterized [6,7], and extensive research has been carried out to elucidate the genetic networks that regulate AF biosynthesis [7]. A series of regulatory factors outside of the AF cluster have been proved to control AF biosynthesis, including the velvet complex [8], transcription factors such as NsdC, NsdD, MeaB, and mtfA [9–11], oxidative stress response-related genes [12,13], and epigenetic modifications including chromatin remodeling and histone acetylation [14,15]. The involvement of these distinct regulatory factors suggests that a multilevel complex regulatory network governs AF biosynthesis.

The conserved heterotrimeric velvet complex, composed of LaeA, VeA, and VelB, is an important regulatory unit in filamentous fungi that couples light-responsive development and secondary metabolism [8]. The nuclear protein LaeA, which contains an *S*-adenosylmethionine (SAM)-binding motif required for its function, is known as a global regulator of secondary metabolism in *Aspergillus* [16]. LaeA is required for the transcriptional activation of *aflR* as well as AF production, and transcriptional profiling of *laeA* mutant strains revealed its global regulatory role in secondary metabolite gene clusters [4,17]. It has been suggested that LaeA controls secondary metabolism epigenetically by altering the chromatin structure to modify the gene expression [18]. Additionally, LaeA was also shown to control developmental differentiation, such as conidiation in various fungi [19–22]. Studies in *Aspergillus nidulans* revealed that LaeA directs the formation of the LaeA-VeA-VelB velvet complex and VelB-VosA complex, the second complex that controls both sexual and asexual development [8,23]. In *A. flavus*, *laeA* deletion mutants displayed loss of sclerotia formation as well as reduction in conidiation, and were less capable of colonizing peanut and maize seeds [17,20].

Polarized growth is one of the distinguishing features of filamentous fungi. In *A. nidulans*, a conidium grows isotropically until first mitosis, and then switches to polarized growth and forms an elongating germ tube, a process that is tightly coordinated with nuclear division [24]. A critical aspect of nuclear division is the interaction of the kinetochore with spindle microtubules. Kinetochore is a multiprotein complex assembled on the centromeres that serves as an interface between chromosomes and spindle microtubules during cell division [25]. The kinetochore functions as a mechanical latch that hooks onto microtubules to support the directional movement of chromosomes and acts as a sensor of proper microtubule attachment, which is coupled with the spindle checkpoint pathway [26,27].

The structure of kinetochore is conserved in eukaryotic species, despite the divergence of its subunits in the primary sequence [28,29]. The Spc105p protein is a crucial member of the kinetochore protein family in *Saccharomyces cerevisiae* and is responsible for correctly binding the kinetochore to spindle microtubules, as well as recruiting the checkpoint proteins [30]. In *Schizosaccharomyces pombe*, the Spc105p ortholog Spc7 has been implicated in spindle integrity and spindle checkpoint signaling [31,32]. The segregation of chromosomes determines the development of *Aspergilli* to a large extent, and a loss of certain genes related to chromosome segregation greatly influences vegetable growth [33,34]. Spc105, the unique ortholog of *S. cerevisiae* Spc105p in *A. flavus*, is a putative chromosome segregation protein and has not yet been investigated to our knowledge.

In this study, we demonstrate for the first time that the conserved kinetochore protein Spc105 in *A. flavus* regulates developmental differentiation, including conidia development and sclerotia production, and secondary metabolism. We further showed that Spc105 interacts directly with the global regulator LaeA, and that a functional interdependence exists between the two proteins. The relationship between Spc105 and LaeA highlights the coordinated interplay between distinct regulators, ensuring precise fungal development and secondary metabolism.

## Results

### Preliminary characterization of *A. flavus* Spc105

The *spc105* ORF (open reading frame) of *A. flavus* consists of 2,318 nucleotides, with two introns, and encodes a putative chromosome segregation protein (Spc105) containing 739 amino acids. The predicted *A. flavus* Spc105 containing 739 amino acids (aa) that harbors a Spc7 (the Spc105p ortholog in *S. pombe*) kinetochore protein domain (residues 195-525 aa), and a Spc7_N domain (residues 1-138 aa) was 26% and 31% identical to *S. cerevisiae* Spc105p and *S. pombe* Spc7, respectively (Fig 1A). A bipartite nuclear localization signal (residues 229-244 aa) and a leucine zipper motif (residues 367-450 aa), which might mediate protein-protein interactions, were predicted by the program PROSITE [35]. The structural analysis of Spc105 proteins from several species showed that all analyzed fungi and bacteria share a conserved Spc7 domain (Fig 1B), and *A. flavus* Spc105 most closely resembles its ortholog in *Penicillium subrubescens* (Fig 1C).

**Fig 1.**
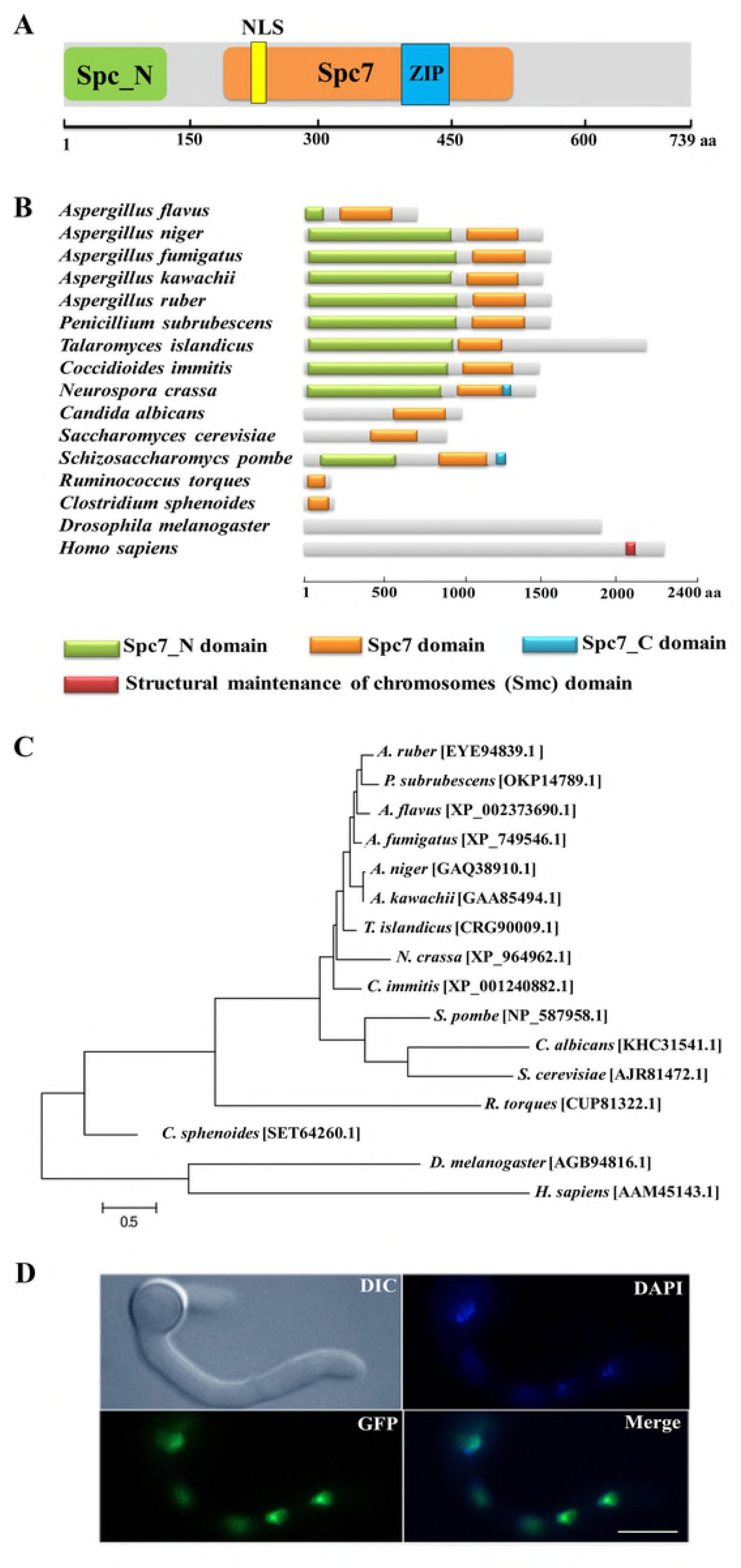
Preliminary characterization of *A. flavus* Spc105. (A) Graphical representation of *A. flavus* Spc105 protein. NLS = bipartite nuclear localization signal, ZIP = leucine zipper motif. (B) Domain analysis of Spc105 protein between different species. (C) Phylogenetic analysis of Spc105 proteins. A neighbor-joining phylogenetic tree was constructed based on sequence alignments of Spc105 proteins using ClustalX2. (D) Localization of Spc105 in *A. flavus*. GFP was fused to the N-terminal of Spc105. Nuclei were stained with 100 ng/mL of 4′,6-diamidino-2-phenylindole (DAPI) and examined by fluorescence microscope.

Through green fluorescent protein (GFP) labeling we found that the Spc105 protein is localized in the nucleus of *A. flavus* during hyphae growth (Fig 1D). Quantitative real-time PCR (qRT-PCR) analysis of *spc105* gene expression in a wild-type (WT) strain showed that it is a low-expression-level gene, and its expression was almost constant during the vegetative growth phase (data not shown), implying that *spc105* is a constitutively expressed gene in *A. flavus*.

### Spc105 influences vegetative growth and conidia germination

To investigate the role of Spc105 in *A. flavus*, we created *spc105* gene deletion (*Δspc105*) and overexpression (*OE::spc105*) mutants (S1 Fig). The *spc105* deletion mutant displayed inhibited colony growth on all tested culture plates including potato dextrose agar (PDA), glucose minimal media (GMM), and Czapek’ media (CZ) (Fig 2 and data not shown). This inhibitory effect was stronger at 25°C than at 37°C compared to the WT and *OE::spc105* strains. Moreover, conidia germination of the *Δspc105* strain was obviously delayed, especially at 25°C (Fig 2). When conidia were inoculated in potato dextrose broth (PDB) medium, *spc105* deletion caused approximately 1 h and 5 h delays in germination at 37°C and 25°C, respectively. There was no conidial swelling for up to 10 h in the *Δspc105* strain at 25°C. After 18 h incubation, conidia of the WT and *OE::spc105* strains had germinated almost 100% while only about 60% of the *Δspc105* conidia had germinated at 25°C. Surprisingly, although the growth and germination rates of the *Δspc105* strain were both delayed, the total mycelia mass in liquid culture was not significantly different from the WT after 24 h incubation (S2 Fig). These results suggest that the *Δspc105* strain is cold-sensitive and Spc105 affects normal hyphal growth and conidia germination in *A. flavus*.

**Fig 2.**
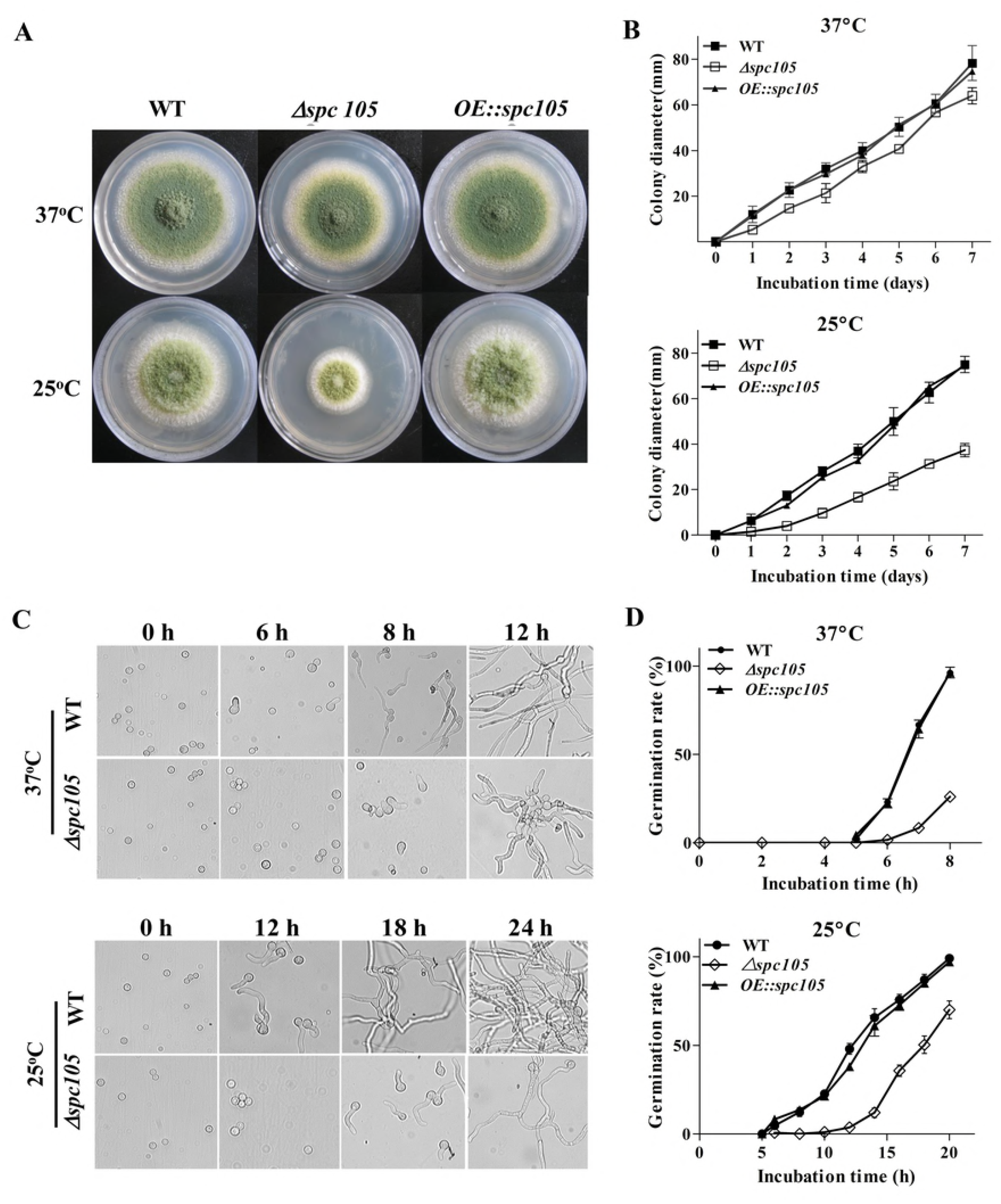
Spc105 affects colony growth and spore germination of *A. flavus*. (A) Colony growth of *spc105* mutants on PDA plates after 5 days incubation. (B) Colony diameter measurement of each strain. (C) Examination of spore germination in PDB culture. (D) Comparison of germination rate of each strain.

### Spc105 regulates conidiophore development and sclerotia production

Microscopic observations revealed that *spc105* deletion resulted in preferential development of conidiophore in *A. flavus*. The *Δspc105* strain produced denser, smaller conidial heads compared with those of WT and *OE::spc105* strains (Fig 3A). Conidiophore stipes were slightly shorter in the *Δspc105* strain, resulting in a somewhat flat colony phenotype in contrast to the typical floccose appearance of WT and *OE::spc105* strains (Fig 3A). Moreover, the *Δspc105* strain produced degenerate conidiophores in submerged culture, a condition which normally completely blocks asexual development [36], while no conidiophores were observed in the WT and *OE::spc105* strains (Fig 3A). Further quantitative analysis of conidial production showed a 2-fold increase in the amount of conidia in the *Δspc105* strain and a slight reduction in the *OE::spc105* strain compared to WT (Fig 3B). qRT-PCR analysis of the conidial development regulatory genes revealed that *spc105* deletion remarkably increased *brlA* transcription levels during vegetative growth in liquid culture (Fig 3D), whereas there was a minimal impact on the transcription of *abaA* and *wetA*.

**Fig 3.**
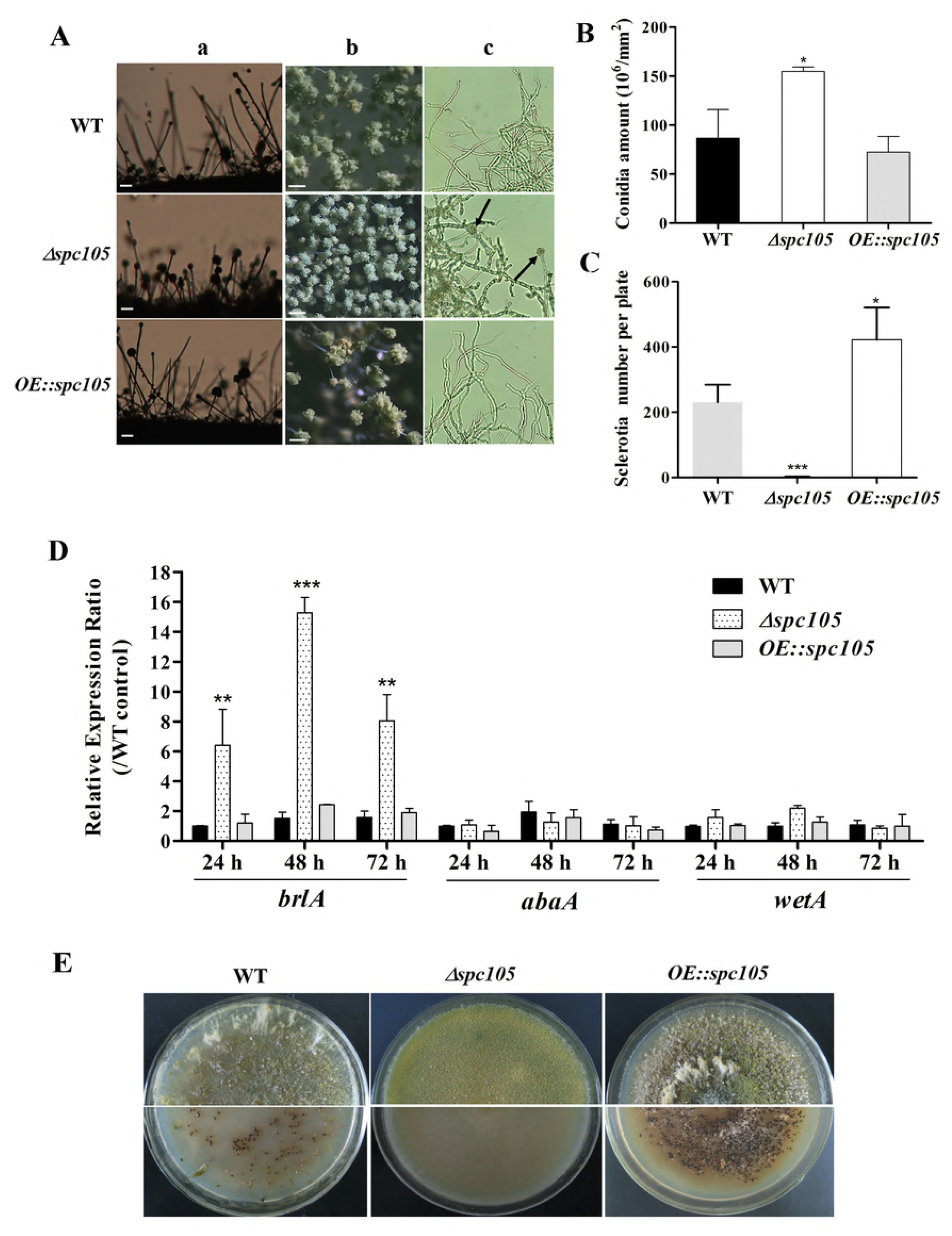
Spc105 regulates conidia development and sclerotia formation in *A. flavus*. (A) Conidia development of each strain cultured at 30°C. a, microscopic observation of conidiophore formation after 24 h incubation on PDA plates. b, examination of conidial head structure under a stereomicroscope after 48 h incubation on PDA plates. Bars represent 200 μm. c, microscopic examination of conidiophore formation after 48 h incubation in submerged PDB culture. Arrows indicate the degenerate conidiophores. (B) Quantitative analysis of conidial production on PDA plates. (C) Statistic analysis of sclerotia production. (D) Relative gene expression of *brlA*, *abaA*, and *wetA*. Strains were cultured in PDB and gene expression levels were normalized (ΔΔCT analysis) to *A. flavus* β-actin gene expression levels. (E) Observation of sclerotia formation. Each strain was grown on CZ agar plates for 2 weeks in the dark at 30°C. \**P* < 0.05; \*\**P* < 0.01; \*\*\**P* < 0.001.

Additionally, *spc105* deletion abolished the production of the resistant structure sclerotia while *spc105* overexpression enhanced sclerotia production (Fig 3E). No sclerotia were formed in the *Δspc105* strain compared with the WT (230 ± 20) and *OE::spc105* strains (422 ± 21) after 14 days of incubation in the dark on sclerotia conducive CZ plates (Fig 3C). The above results suggest that Spc105 regulates conidia development and sclerotia formation in *A. flavus*.

### Deletion of *spc105* delays the nuclear division cycle

DAPI staining of nuclei during conidia germination revealed that *spc105* deletion delayed the nuclear division cycle in *A. flavus* (Fig 4). When cultured in PDB at 37°C, the majority (approximately 70%) of WT and *OE::spc105* conidia had undergone the first round of nuclear division within 4 h after inoculation. The conidial population of the WT strain was 25% uninucleate, 70% binucleate, and 5% trinucleate 4 h after inoculation. In contrast, conidia of the *Δspc105* strain were 80% uninucleate at 4 h, and they underwent their first nuclear division at 6 h (Fig 4A). At 25°C, the number of nuclei increased slowly as a function of time. Up to 8 h, approximately 70% of WT and *OE::spc105* conidia had 2 nuclei and 10% had 4 nuclei, while 65% of the *Δspc105* conidia were still uninucleate (Fig 4A). Fig 4B shows the mean number of nuclei per germling at each time point. The results indicated that *spc105* deletion in *A. flavus* caused nuclei to transiently arrest in interphase, although nuclear division was eventually completed in the *Δspc105* strain. Further examination of the expression of several mitosis-related genes [37] revealed that the expression levels of *cdc7* (cell division cycle 7), *cdk* (cyclin-dependent kinase A), and three (*mcm2, mcm3* and *mcm6*) out of the six *mcm* (minichromosome maintenance proteins 2-7) genes were downregulated in the absence of *spc105*. Alternatively, the expression of *nimE* (cyclin B) and *nimO* (dbf4) did not show significant differences (Fig 4C). These results confirmed that Spc105 is involved in the nuclear division of *A. flavus*.

**Fig 4.**
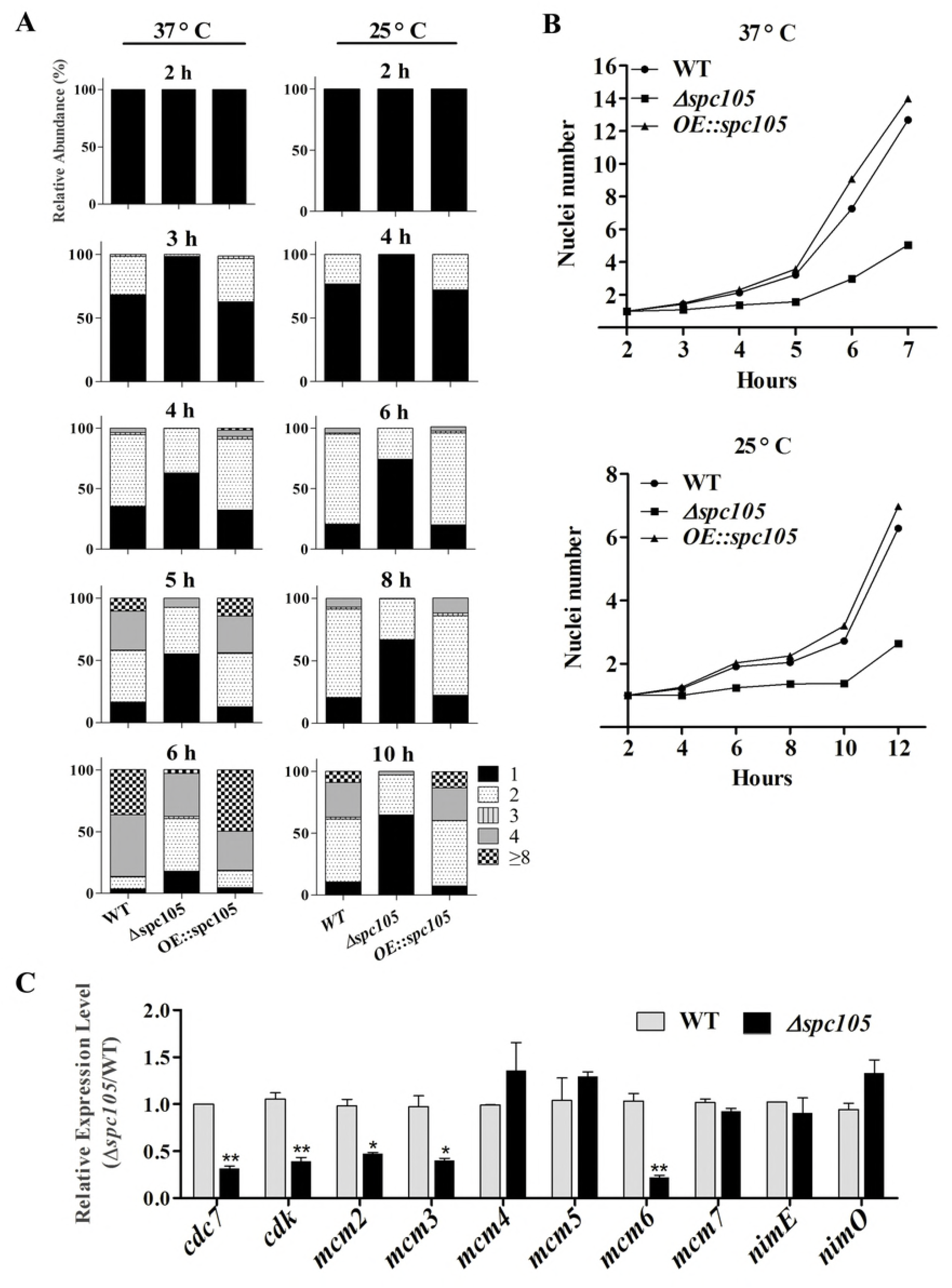
Involvement of Spc105 in the nuclear division of *A. flavus*. (A) Graphic image of the counted nuclei, the numbers 1, 2, 3, 4 and ≥ 8 represent the nuclei number. (B) The mean number of nuclei per germling of each strain. (C) qRT-PCR analysis of meiosis-related gene expression. Strains were incubated for 20 h in 30 mL PDB at 25°C and mycelia samples were collected for RNA extraction. Relative expression ratios were calculated relative to the WT control. Error bars represent the standard deviation based on three replicates.

### Spc105 affects secondary metabolite production and colonization of *A. flavus*

*A. flavus* contains 56 secondary metabolite clusters in its genome and produces a wide variety of secondary metabolites, the biosynthesis of which is often associated with morphological differentiation [38]. Aflatoxin B_1_ (AFB_1_) is the most crucial and abundant metabolite in *A. flavus*. Detection of both the culture and mycelia extracts showed that *spc105* deletion almost completely abolished AFB_1_ production while *spc105* overexpression resulted in increased AFB_1_ levels (Fig 5A), suggesting a significant positive effect of Spc105 on AF production. The production of kojic acid, an important chemical material, was also positively affected by Spc105 (Fig 5B). In addition, the production of the mycotoxin CPA was increased in the *Δspc105* strain and slightly reduced in the *OE::spc105* strain, compared with WT (Fig 5C). The above results were confirmed by high-performance liquid chromatography (HPLC) analysis (S3 Fig). All analyses indicated that Spc105 exerts a role in *A. flavus* secondary metabolite biosynthesis.

**Fig 5.**
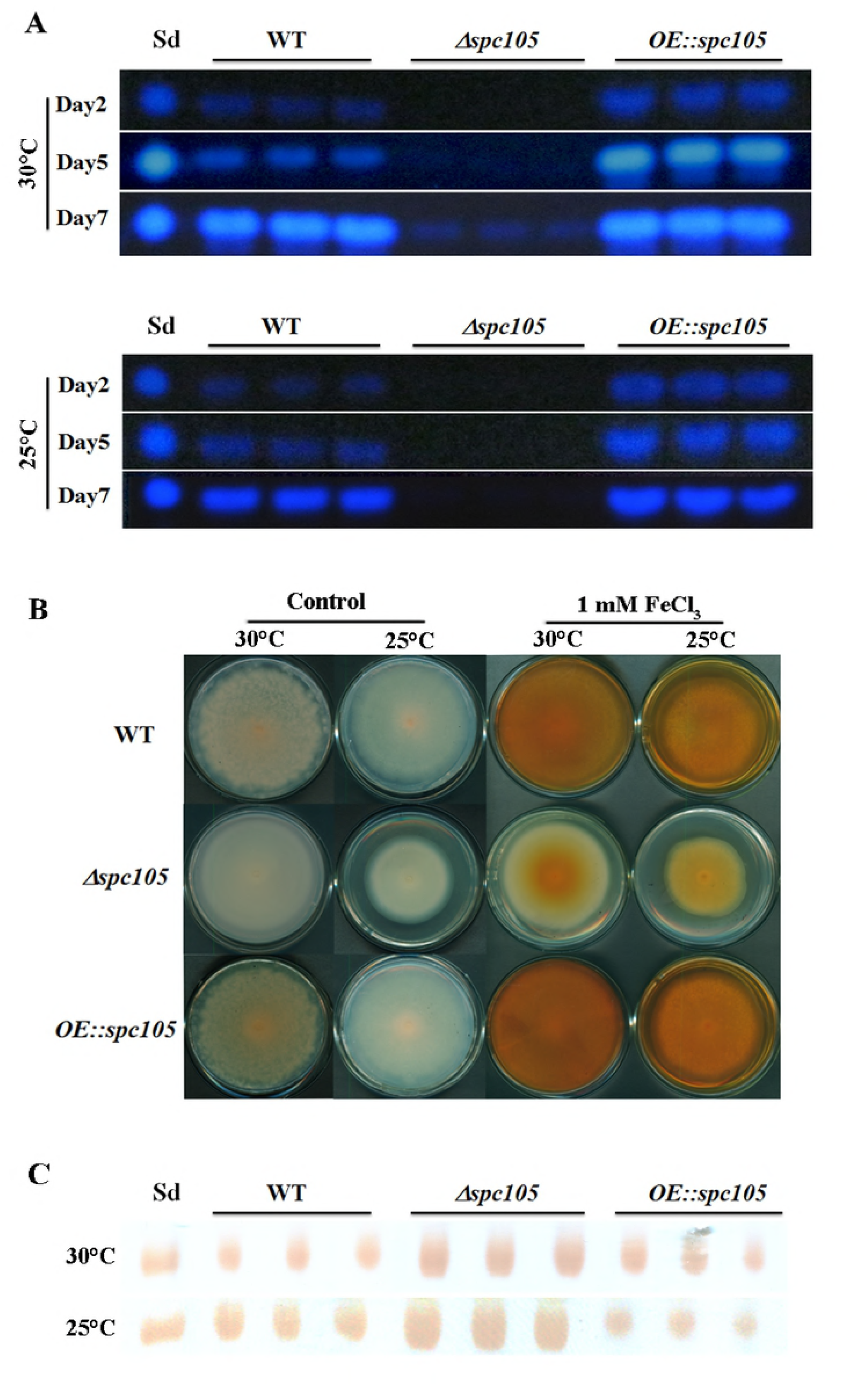
Spc105 regulates the production of aflatoxin, kojic acid, and cyclopiazonic acid. (A) Time course semi-quantitative thin-layer chromatography (TLC) analyses of aflatoxin B_1_ (AFB_1_) production in the PDB culture. Sd represents the AFB_1_ standard. (B) Determination of kojic acid production in solid medium through the colorimetric method. Strains were incubated on PDA plates for 7 days. Images are representative of four experimental replicates. (C) TLC detection of CPA production in PDB liquid culture of each strain. Sd represents the CPA standard.

In addition, surface-sterilized peanut seeds were inoculated with the above three *A. flavus* strains to assess the host colonization capability of *spc105* mutants. Results showed that the *Δspc105* strain grew less vigorously on peanuts than WT and *OE::spc105*, corresponding to a significant decrease in the conidial production on seeds (Fig 6A and Fig 6B). Moreover, AFB_1_ was not detected in the *Δspc105* infected seeds (Fig 6C).

**Fig 6.**
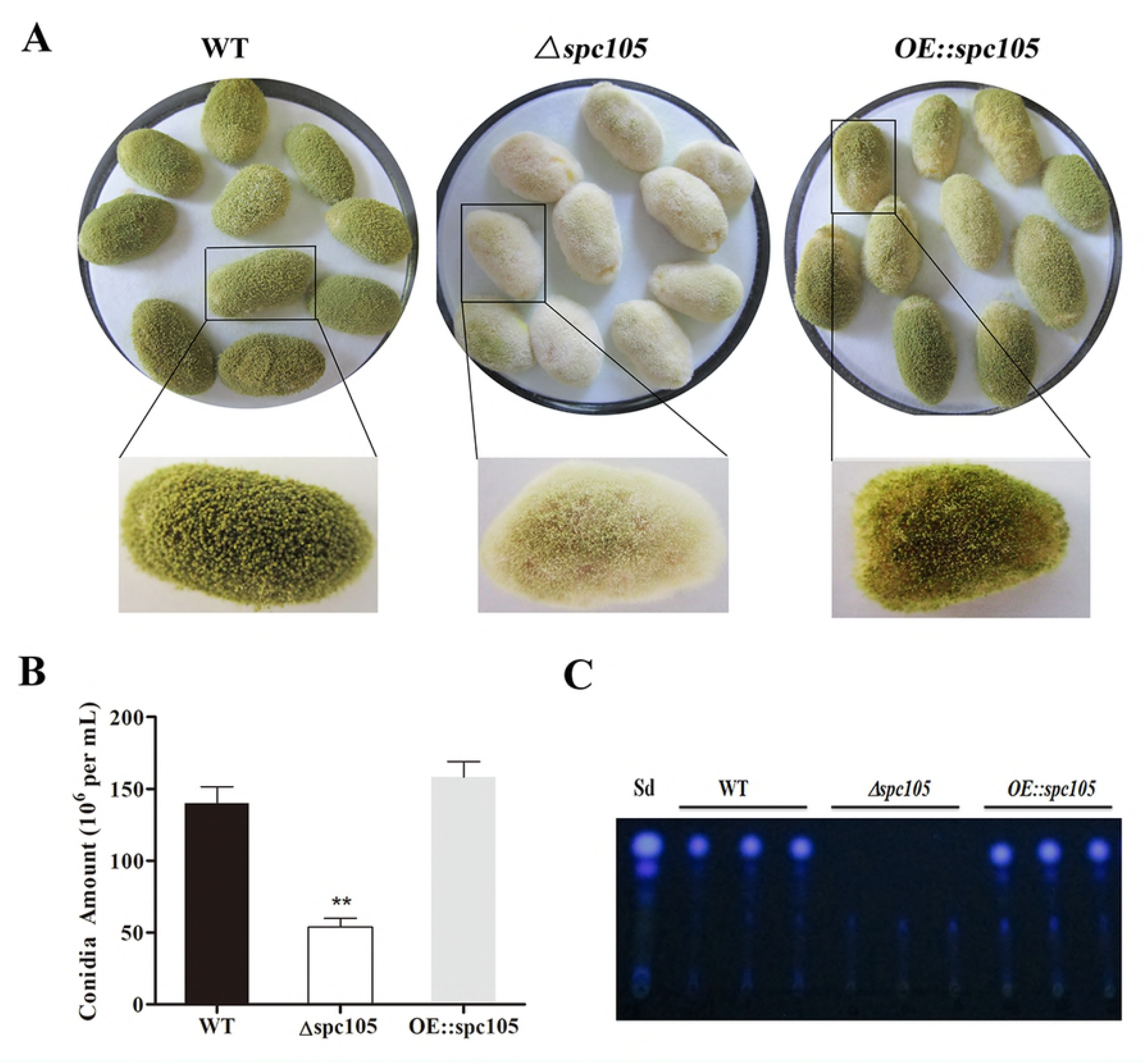
Growth and aflatoxin production of WT, *Δspc105*, and *OE::spc105* strains on peanut seeds. (A) Photographs of peanut seeds infected with *A. flavus* strains after three days of incubation at 30°C. (B) Quantification of conidia from the infected peanut seeds by strain. \*\**P* < 0.01. (C) TLC analysis of AFB_1_ levels in infected peanuts.

### Transcriptome comparison between *Δspc105* and WT

Differential gene expression analysis, carried out by assessment of the genome wide transcriptional profile between *Δspc105* and WT strains cultured at 25°C for 48 h, identified 1,846 differentially expressed genes (DEGs), including 670 upregulated and 1,176 downregulated genes (S4 Fig).

Functional analysis based on Gene Ontology (GO) and Kyoto Encyclopedia of Genes and Genomes (KEGG) pathway annotation revealed that the downregulated genes were involved in various cellular processes mainly associated with AF biosynthesis, fatty acid metabolism, rRNA processing, and the ribosome biogenesis-related process (S5 Fig). The significant downregulation of AF biosynthetic genes was consistent with the above AF detection results. Further, the upregulated gene set was mainly implicated in the process of carbohydrate metabolism, cell proliferation, protein phosphorylation, and ubiquitination. These results indicated that the absence of *spc105* had an extensive and complex effect on cell division, primary and secondary metabolism, and protein modification in *A. flavus*.

By focusing on the transcriptional changes in secondary metabolism genes induced by *spc105* deletion, we found that 23 out of the 74 backbone genes were differentially expressed, corresponding to 19 out of the predicted 56 secondary metabolite gene clusters, including the AF cluster (# 54), CPA cluster (# 55), and kojic acid cluster (# 56) (S1 Table). Among the 34 AF cluster genes, 31 genes were significantly downregulated in the *Δspc105* strain, including the AF cluster-specific regulatory genes *aflR* and *aflS* (Fig 7). These expression profiles were verified by qRT-PCR analysis (S2 Table).

**Fig 7.**
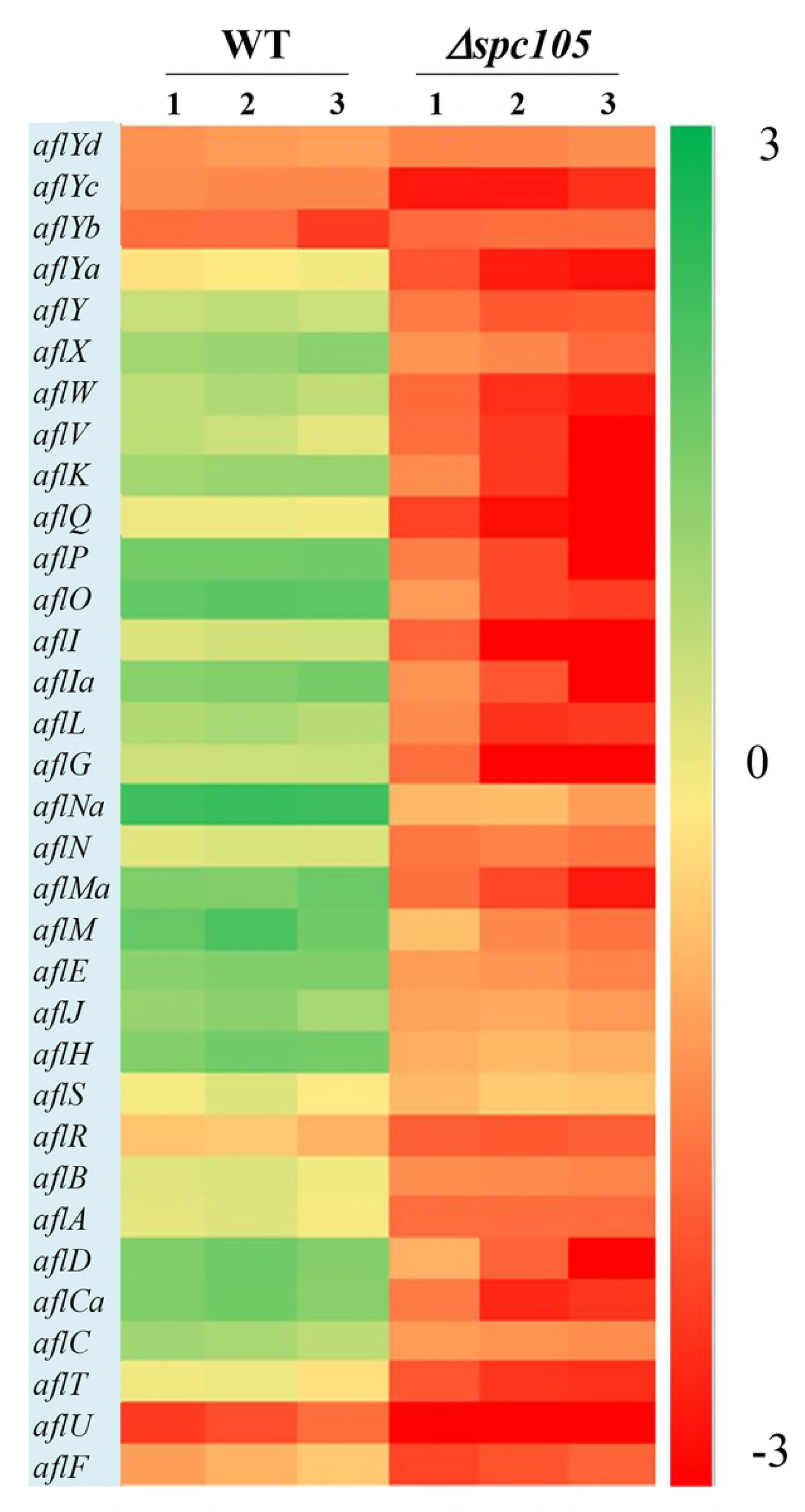
Gene expression analysis of AF cluster genes. The heatmap shows gene expression levels in *Δspcl05* and WT samples. Gene expression is plotted with colors corresponding to the log10 value of the FPKM values of each gene.

In addition to the annotated AF cluster genes, several upstream regulatory genes were differentially expressed in our data, including the velvet complex encoding genes *laeA*, *velB*, and *velC* (another velvet family member) and the signal transduction related genes *pkaC* and *rasA* [39] (Table 1). These results may explain the observed phenotype and secondary metabolite production changes in the *Δspc105* background.

**Table 1.**
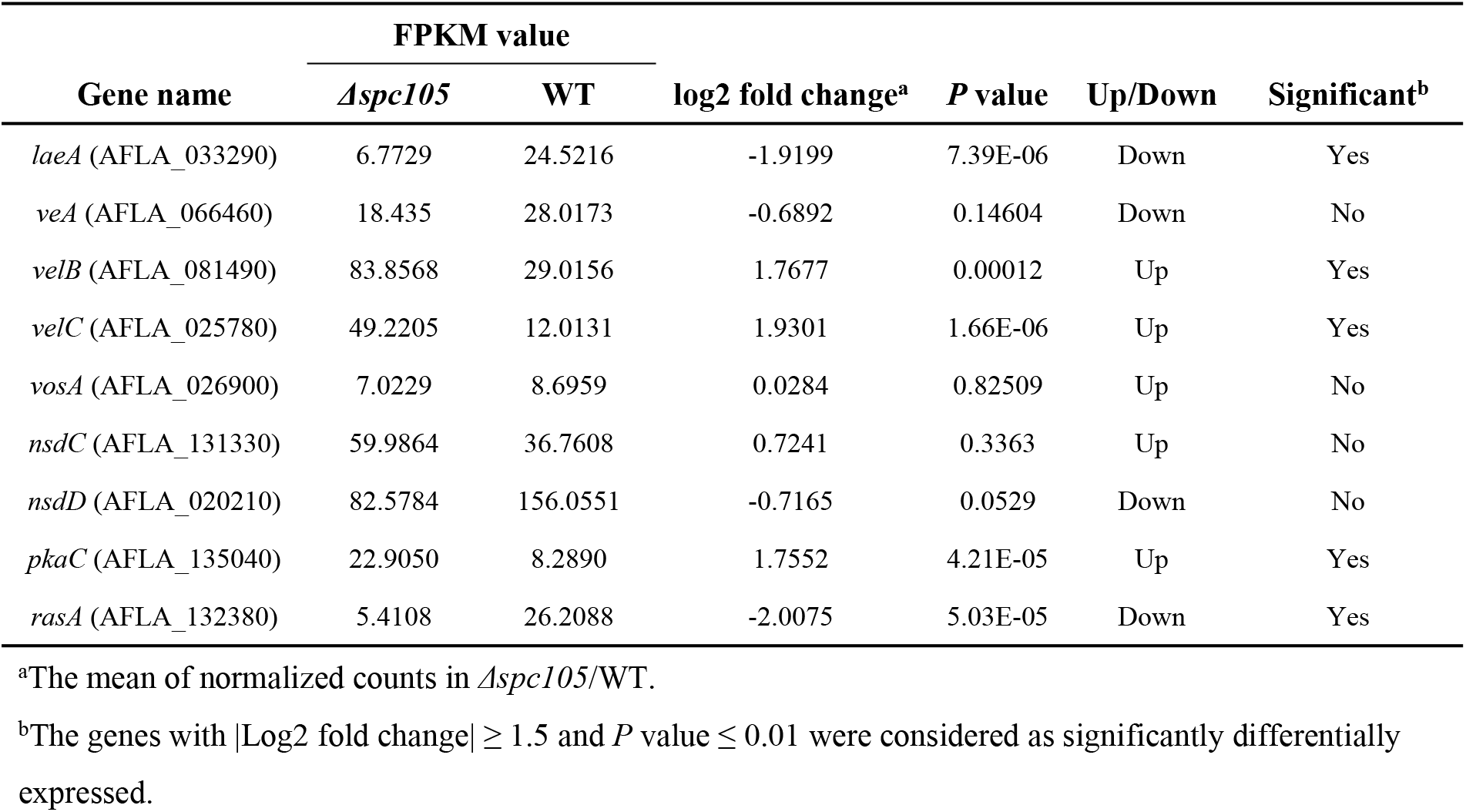
Effects of *spc105* deletion on regulatory gene expression in *A. flavus*

| Gene name | FPKM value |  | log2 fold change<sup>a</sup> | P value | Up/Down | Significant<sup>b</sup> |
| --- | --- | --- | --- | --- | --- | --- |
|  | *Δspc105* | WT |  |  |  |  |
| *laeA* (AFLA_033290) | 6.7729 | 24.5216 | -1.9199 | 7.39E-06 | Down | Yes |
| *veA* (AFLA_066460) | 18.435 | 28.0173 | -0.6892 | 0.14604 | Down | No |
| *velB* (AFLA_081490) | 83.8568 | 29.0156 | 1.7677 | 0.00012 | Up | Yes |
| *velC* (AFLA_025780) | 49.2205 | 12.0131 | 1.9301 | 1.66E-06 | Up | Yes |
| *vosA* (AFLA_026900) | 7.0229 | 8.6959 | 0.0284 | 0.82509 | Up | No |
| *nsdC* (AFLA_131330) | 59.9864 | 36.7608 | 0.7241 | 0.3363 | Up | No |
| *nsdD* (AFLA_020210) | 82.5784 | 156.0551 | -0.7165 | 0.0529 | Down | No |
| *pkaC* (AFLA_135040) | 22.9050 | 8.2890 | 1.7552 | 4.21E-05 | Up | Yes |
| *rasA* (AFLA_132380) | 5.4108 | 26.2088 | -2.0075 | 5.03E-05 | Down | Yes |

<sup>a</sup>The mean of normalized counts in *Δspc105*/WT.
<sup>b</sup>The genes with |Log2 fold change| $\geq 1.5$ and *P* value $\leq 0.01$ were considered as significantly differentially expressed.

### Interdependent relationship between Spc105 and LaeA

Since *spc105* deletion reduced the expression of the global regulatory gene *laeA*, we hypothesized that Spc105 may correlate with LaeA to regulate secondary metabolism. To ascertain the potential functional relationship between Spc105 and LaeA, we generated *Δspc-OElaeA* and *OEspc-ΔlaeA* double mutant strains (S1 Fig). Results showed that fungal development, AF production, and AF cluster gene expression were similar between the *Δspc-OElaeA* and *Δspc105* strains (Fig 8 and Table 2), indicating that overexpression of *laeA* cannot restore the development and the AF production defects of the *Δspc105* strain. Likewise, the *OEspc-ΔlaeA* mutant exhibited properties like those of *ΔlaeA*, suggesting that *spc105* overexpression failed to rescue the effects caused by the lack of *laeA* in *A. flavus*. We also observed that all above mutants displayed reduced expression levels of AF cluster genes, including *aflR* and *aflS*. These results suggested that Spc105 and LaeA hold interdependent functions. In addition, the transcription level of *spc105* was elevated in the *ΔlaeA* background, where it was slightly decreased in the *OE::laeA* strain (Fig 8C), suggesting a negative regulatory effect of *laeA* on *spc105* gene transcription.

**Fig 8.**
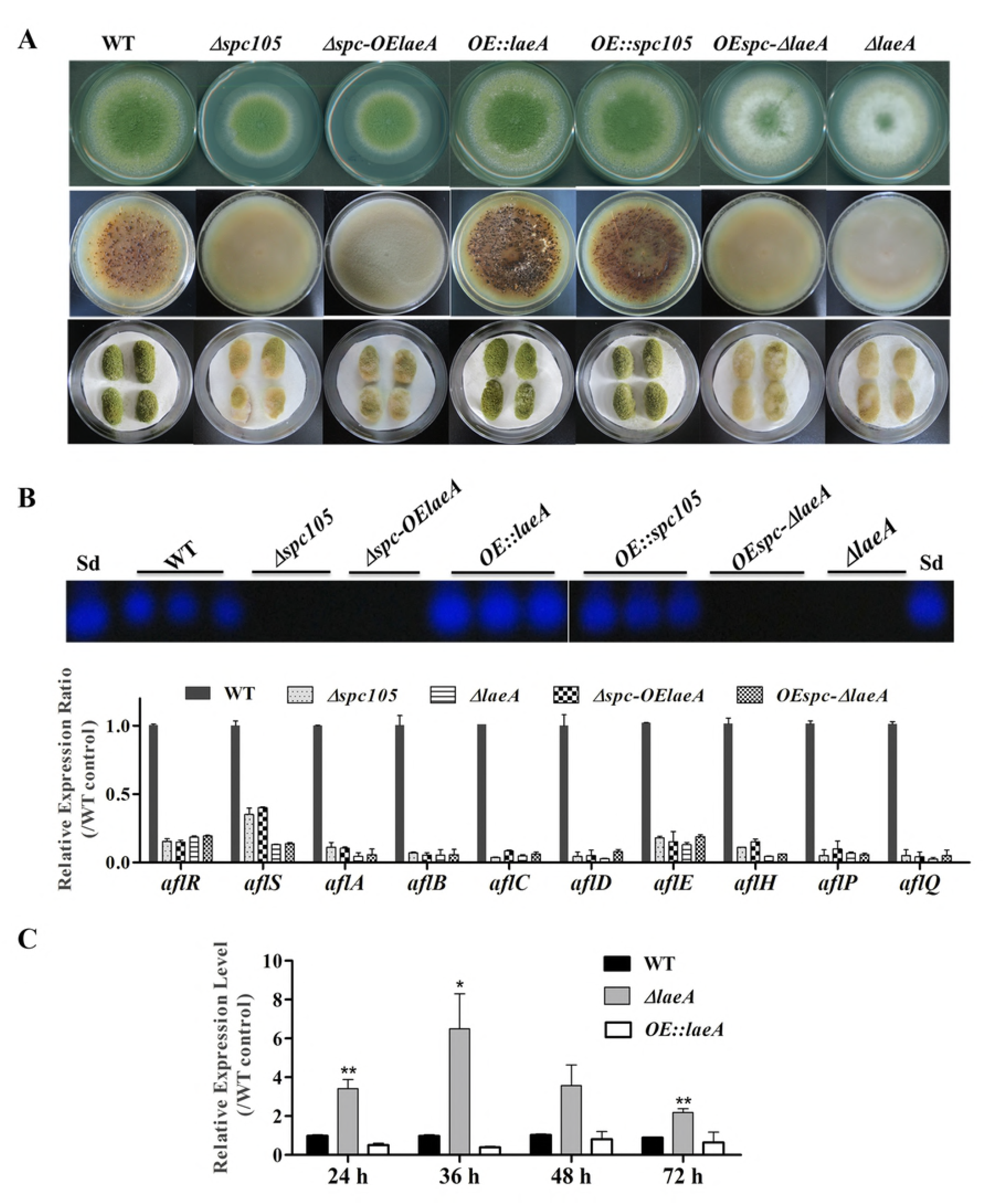
Relationship between Spc105 and LaeA. (A) Phenotypes of *spc105/laeA* double mutant strains. Upper: Phenotypes of mutant strains after 5 days incubation at 30°C on PDA plates. Middle: Sclerotia production of each strain on CZ plates after 2 weeks incubation in the dark. Bottom: Peanut infection of each strain after 3 days incubation at 30°C. (B) TLC analyses of AFB_1_ production of each strain and gene expression analysis of AF cluster genes in the PDB culture after 48 h incubation at 30°C. (C) Effect of *laeA* deletion and overexpression on *spc105* gene expression. Mycelia were harvested from PDB culture at the indicated time point. Error bars represent the standard deviations based on three replicates.

**Table 2.** Summary of *A. flavus spc105/laeA* double mutant phenotypes^a^

| Parameters | WT | *Δspc105* | *Δspc105-OElaeA* | *OE::laeA* | *OE::spc105* | *OEspc105-ΔlaeA* | *ΔlaeA* |
| --- | --- | --- | --- | --- | --- | --- | --- |
| Radial growth<sup>b</sup> | 51 ± 1.5 | 35 ± 1.5 | 36 ± 1.2 | 51 ± 0.4 | 52 ± 1.0 | 51 ± 0.8 | 50 ± 1.2 |
| Germination rate<sup>c</sup> | 100% | 50% | 51% | 100% | 100% | 100% | 100% |
| Conidia production<sup>d</sup> | 86 x 10<sup>6</sup> | 155 x 10<sup>6</sup> | 148 x 10<sup>6</sup> | 75 x 10<sup>6</sup> | 70 x 10<sup>6</sup> | 45 x 10<sup>6</sup> | 42 x 10<sup>6</sup> |
| Sclerotia production | 230 ± 20 | ND<sup>e</sup> | ND | 505 ± 10 | 422 ± 21 | ND | ND |
| Degenerate conidiophores formation in submerged culture | ND | Yes | Yes | ND | ND | ND | ND |
| Peanuts infection | Control | Reduced | Reduced | Enhanced | Enhanced | Reduced | Reduced |

<sup>a</sup>All strains were incubated at 30°C for phenotype observation. All analyses were performed in triplicate.
<sup>b</sup>Radial growth on PDA plates after 5 days incubation, expressed in mm.
<sup>c</sup>Germination rate was recorded after 10 h incubation in PDB culture when WT spores were 100% germinated.
<sup>d</sup>Conidia amount per mm<sup>2</sup> on PDA plates was recorded.
<sup>e</sup>ND: not detected

### Spc105 interacts directly with LaeA

The finding of functional interdependence between Spc105 and LaeA led us to hypothesize that Spc105 could directly interact with LaeA. To test this hypothesis, a yeast two-hybrid (Y2H) assay was performed and the results showed that the yeast cells co-transformed with AD-Spc105 and BD-LaeA plasmids exhibited positive galactosidase activity (Fig 9A). This result indicates that Spc105 directly interacts with LaeA. Similar results were obtained in yeast cells co-transformed with BD-Spc105 and AD-LaeA plasmids. The Spc105-LaeA interaction was also confirmed by the *in vitro* GST pull-down assay with recombinantly expressed GST-Spc105 and His_6_-LaeA (Fig 9B). Additionally, we observed negative results in examining whether Spc105 interacts with VelB or VeA, as determined by Y2H assay (data not shown).

**Fig 9.**
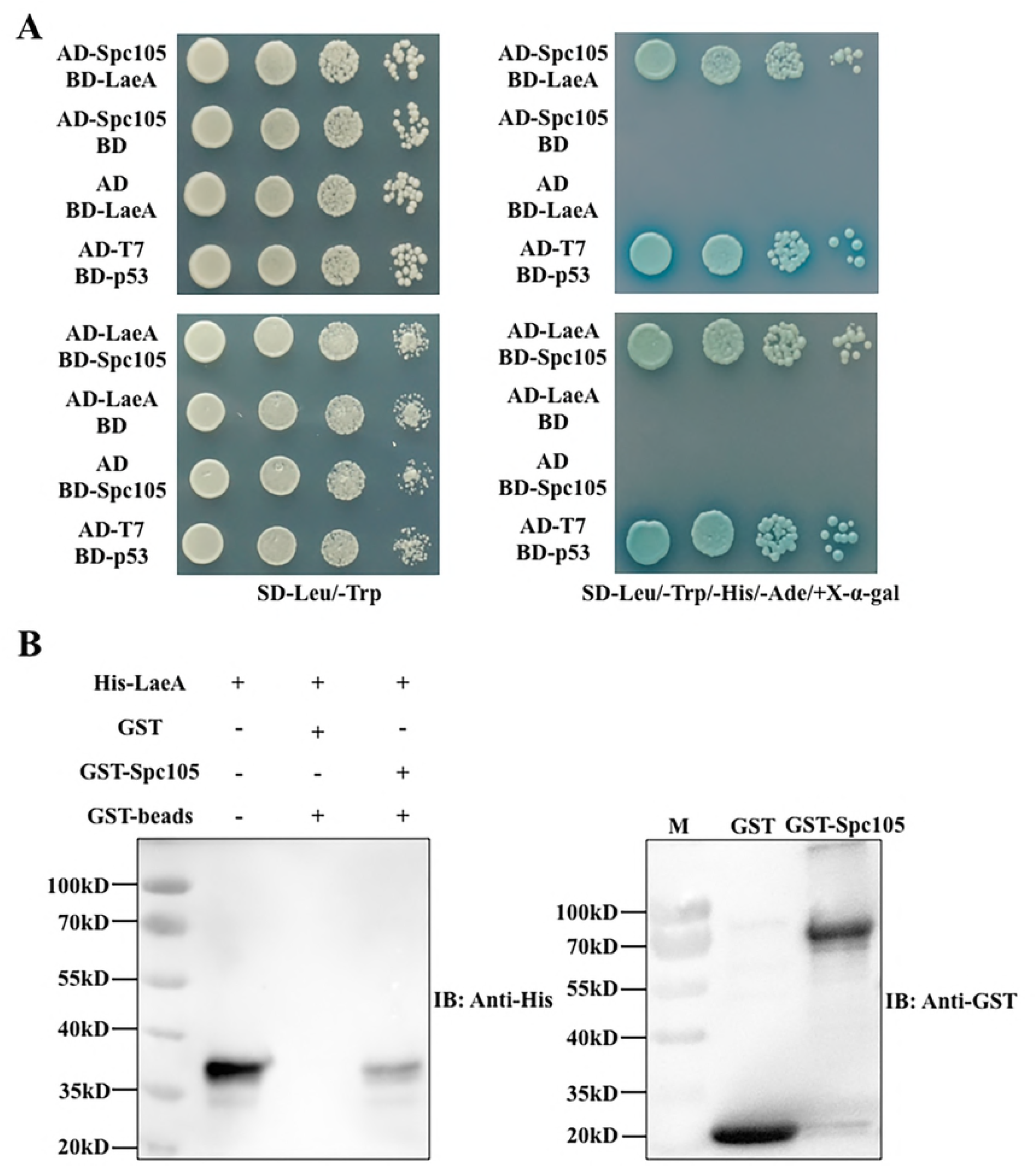
Spc105 interacts with LaeA in the Y2H and GST pull-down assays. (A) Y2H assays to determine protein-protein interactions. Yeast cells were grown in liquid selective medium overnight and diluted serially. Four microliters of serially diluted yeast cells were spotted on selective synthetic dropout media SD/-Leu/-Trp/-His/-Ade/+X-α-gal and incubated at 30°C for 3-5 days. The SD/-Leu/-Trp plate is nonselective and served as the loading control. (B) GST pull-down assay of the interaction between Spc105 and LaeA *in vitro*. Recombinant GST and GST-Spc105 were incubated with recombinant His_6_-LaeA and subsequently purified by glutathione magnetic beads. Immunoblot analysis was performed to detect the presence of His_6_-LaeA using an anti-His-tag antibody.

LaeA is a constitutively nuclear protein and its interaction with VeA-VelB in the nucleus is required for the control of secondary metabolism and development [8,23]. Since both VeA and Spc105 can interact with LaeA, we investigated the interaction domains of LaeA-Spc105 and LaeA-VeA. Y2H assay of the interactions between full length LaeA with 6 different truncations of Spc105 revealed that Spc105-LaeA interaction depends on the Spc7 domain (203-526 aa), and that the leucine zipper (367-450 aa) in the Spc7 domain is required for this interaction (Fig 10A). Further, 5 different truncated forms of LaeA were tested as interaction partners with full length Spc105 and VeA. Our results showed that Spc105 can interact with the SAM binding domain (128-285 aa) of LaeA, whereas VeA can only interact with full length LaeA (Fig 10B). These results suggest that there may be a competition between the nuclear Spc105 and the light-sensitive protein VeA, which shuttles between the cytoplasm and nucleus.

**Fig 10.**
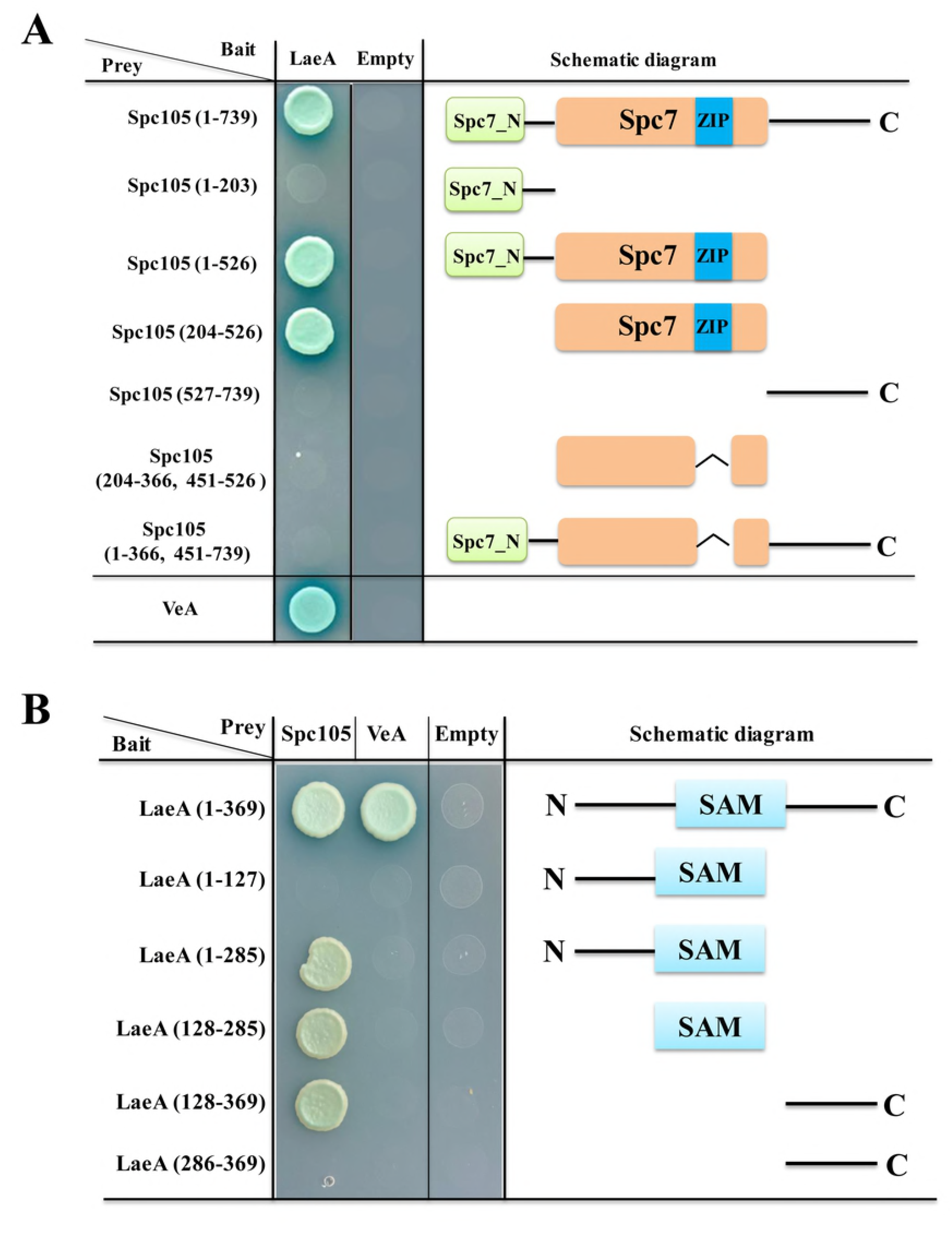
Interaction domain mapping of Spc105 and LaeA. (A) Six truncation mutants of Spc105 were constructed and tested against full length LaeA in the yeast-two-hybrid assay. (B) Five different truncations of LaeA were tested as interaction partners with full-length Spc105 and VeA.

## Discussion

Accurate chromosome segregation is crucial for cellular and organismal viability and requires coordinated interaction between the spindle and kinetochore [27]. As a key member of the kinetochore, the participation of Spc105 in the kinetochore-spindle interaction and the spindle assembly checkpoint (SAC) during chromosome segregation has been widely studied in model organisms such as *Drosophila melanogaster*, *Caenorhabditis elegans*, and *S. cerevisiae* [29,40]. However, the role of the Spc105 ortholog in filamentous fungi has not been explored. In this study, we demonstrated that Spc105 interacts directly with LaeA, a global regulator of secondary metabolism, and plays multiple roles in developmental differentiation and secondary metabolism in *A. flavus*. To the best of our knowledge, this is the first report describing the function of the Spc105 ortholog in filamentous fungi, particularly in secondary metabolism.

Spc105 is functionally conserved in eukaryotes from yeast to humans. In human cells, a lack of hSPC105 activity results in lagging sister chromatids during metaphase and widespread chromosome mis-segregation at anaphase [41]. In *D. melanogaster*, Spc105 inactivation completely abolishes the kinetochore-microtubule attachment [42]. In *C. elegans*, the RNAi-mediated depletion of the Spc105p ortholog KNL-1 prevents kinetochore assembly and leads to a kinetochore-null phenotype [43]. In *S. pombe*, the Spc105p ortholog Spc7 is required for kinetochore attachment and spindle integrity [31]. In the filamentous fungus *A. flavus*, we found that the Spc105 protein harbors a conserved Spc7 domain, and *spc105* gene deletion caused a delay in the nuclear division cycle (Figs 1 and 4). Taken together, these results illustrate that Spc105 holds a conserved role in mitosis in various organisms. The mitotic delay caused by the loss of *spc105* may be due to the activation of a cell cycle checkpoint associated with DNA replication or spindle assembly, allowing time for repair mechanisms to resolve the detected problem. Several genes involved in the cell cycle were differentially expressed by *spc105* gene deletion (Fig 4C). Cdc7 and CDK are conserved kinases that are required to activate the origins of DNA replication throughout the synthesis phase (S phase) [44–46]. During the S phase, the MCM2-7 complex, a heterohexamer of six related proteins that constitutes the replicative DNA helicase, is activated in a Cdc7- and CDK-dependent manner [45]. In our results, decreased expression levels of the *cdc7* and *cdk* genes and the *mcm* family member genes indicated that *spc105* deletion caused a DNA synthesis defect in the S phase.

Emerging data in both yeast and humans link the cell cycle to ribosome synthesis [47,48]. A delay in the cell cycle has been shown to cause a “slow growth signature” and reduced expression of genes associated with ribosome synthesis, while increasing the transcription of genes connected to respiratory growth [48,49]. In our study, the restricted colony growth and delayed spore germination caused by *spc105* deletion, as well as the downregulated ribosome biosynthesis-related gene expression pattern in our RNA-seq data, also support this view (Fig 2 and S5 Fig). The *Δspc105* mutant exhibited varying degrees of impact on colony growth and spore germination at different temperatures, suggesting that the effect on vegetative growth established by *spc105* deletion is dependent on temperature. Furthermore, our study also found that Spc105 regulates conidiation and sclerotia production in *A. flavus* (Fig 3). During growth in PDB submerged culture, *Δspc105* mutants produced degenerate conidiophores (Fig 3A). This phenomenon resembles the reported abnormal conidiophore development observed under submerged cultures in the *nsdC* and *nsdD* deletion strains in *A. flavus*, both of which were found to negatively regulate conidiation [9,36]. The significantly increased expression level of the asexual development transcriptional activator gene *brlA* in *Δspc105* suggests a negative regulatory role of Spc105 in conidia development.

In addition to regulating morphological differentiation, Spc105 also had a broad effect on the secondary metabolism of *A. flavus*. Our transcriptome analysis revealed that *spc105* deletion stimulated transcriptional changes in numerous secondary metabolite cluster genes (S1 Table). Most of these genes were downregulated by *spc105* deletion, including the AF gene cluster (# 54). *Δspc105* and *OE::spc105* strains yielded inhibited and increased AF production, respectively, suggesting a positive regulation of Spc105 on AF synthesis. The loss of AF production in *Δspc105* coincided with a markedly reduced expression of AF cluster genes. This may be due to the decreased transcripts of *aflR* and *aflS* (Fig 7 and S2 Table), encoding transcription factors that act as AF cluster activators [50]. In addition to AF, the biosynthesis of kojic acid, a scavenger of free radicals [51], was also positively regulated by Spc105 (Fig 5). The production of both AF and kojic acid are postulated to relieve oxidative stress [52,53]. In fact, several oxidative stress response-related genes were differentially expressed in the *Δspc105* strain (S3 Table), and further studies on the relationship between Spc105 and oxidative stress response is underway.

In contrast to the effects on AF and kojic acid production, Spc105 exhibits a minimal negative effect on the production of CPA, another mycotoxin produced by various species of *Aspergillus* and *Penicillium*. The deletion of *spc105* elevated the amount of CPA, whereas overexpression of *spc105* yielded the opposite effect (Fig 5). The CPA cluster (# 55) was speculated to be physically linked to the AF gene cluster (# 54), which resides in a subtelomeric region on chromosome 3 of *A. flavus* [54]. Although the reduced expression of 4 genes in the CPA cluster in our *Δspc105* transcriptome data is inconsistent with the increased CPA production, this might be attributed to the different culture times for gene expression detection and CPA production.

In *A. flavus*, the global regulator of secondary metabolism LaeA is required for the transcription of the AF cluster, including the pathway activator gene *aflR* [16,51]. Previous studies indicated that the deletion of *laeA* in *A. flavus* led to loss of sclerotia and AF production, as well as reduced pathogenicity, while overexpression of *laeA* yielded opposite phenotypes [17,20,55] similar to those displayed in the *Δspc105* mutant in our experiments (Fig 8). The reduced *laeA* expression in the *Δspc105* strain shown in the transcriptome data indicated a correlation between Spc105 and LaeA. We further identified the direct interaction between Spc105 and LaeA (Fig 9) and proved that the two proteins share interdependent functions in the control of AF biosynthesis (Fig 8). LaeA is a putative methyltransferase that contains a SAM domain required for its function [16]. The primary role of LaeA is to regulate secondary metabolic gene clusters, and one of the proposed mechanisms for its regulation is that LaeA differentially methylates histone proteins and then alters chromatin structure to modulate gene expression [56]. However, the exact mechanism of how LaeA regulates secondary mechanisms remains enigmatic. Our findings showing that Spc105 can interact with the SAM binding domain of LaeA may provide new clues for interpreting the molecular functions of LaeA. Further studies will examine whether Spc105 correlates with chromatin remodeling and how LaeA is involved in this process.

Previous studies in *A. nidulans* showed that the light-sensitive protein VeA bridges VelB and LaeA to form the nuclear velvet complex in the dark, and is required for LaeA to control secondary metabolism [8,16]. Since both LaeA and Spc105 are located in the nucleus while VeA shuttles between the cytoplasm and nucleus, we speculated that Spc105 may compete with VeA for binding to LaeA. Our hypothesis was supported by mapping of the interaction domains of VeA-LaeA and Spc105-LaeA (Fig 10). Additionally, it is also important to note the induced expression of the velvet family members *velB* and *velC* in the *Δspc105* background (Table 1). In *A. nidulans*, the VeA-VelB dimer functions to activate sexual development, and VelB also forms a complex with VosA to repress asexual conidiation in the dark [8,23]. In *A. flavus*, VelB in concert with VeA has a positive effect on conidiation [57]. Thus, the transcriptional change of *velB* might be partially responsible for the observed promoted conidiation in the *Δspc105* strain. Further research is needed to investigate the relationship between Spc105 and the velvet proteins.

The regulation of fungal development and secondary metabolism requires a coordinated interplay of regulators, and the biological properties of a protein depend on its physical interaction with other molecules in the extensive and complex networks of the cell. The Spc105-LaeA identified in our study represents a communication between cell cycle progression and the velvet complex, which may improve our understanding of LaeA-involved regulatory networks. LaeA, in association with the velvet family proteins, was proposed to be involved in supporting the development of progeny in *Aspergillus* by controlling the production of secondary metabolites, which is considered beneficial for fungal survival in their ecological niche [23,58]. From this perspective, we hold the opinion that LaeA functions in concert with Spc105 to improve fungal survival capability. It will be interesting to see whether this interaction is conserved in other fungi and whether LaeA can form complexes with other cell cycle related proteins.

## Material and methods

### Culture conditions

Fungal strains used in this study are listed in S4 Table. All strains were maintained as glycerol stocks and were grown on potato dextrose agar (PDA, Difco) for spore collection at 30°C.

### Strain construction

*A. flavus* NRRL3357 (kindly provided by Gary Payne) was used as the wild-type strain, and *A. flavus* NRRL3357-5 [59], was used for gene disruption or overexpression. All primers used for strain construction are listed in S5 Table. Briefly, the 5′ and 3′ flank regions of *A. flavus spc105* were amplified and fused with the *Aspergillus fumigatus pyrG* marker gene through double joint PCR to generate the *spc105* deletion construct (S1 Fig). To construct the *spc105* overexpression strain, the *A. nidulans gpdA* promoter was fused upstream of the *spc105* coding region, after which the *spc105 5′* flank, *A. fumigatus pyrG*, and *gpdA-spc105* fragments were joined together. To determine the localization of Spc105 in *A. flavus*, an eGFP tag amplified from the pEGFP-C1 vector was inserted at the N-terminus of the Spc105 protein, and driven by the *gpdA* promoter with a five Gly-plus-Ala repeat (GA-5) linker [60]. The above PCR constructs were transformed individually into *A. flavus* NRRL3357-5 protoplasts based on the polyethylene glycol method [61]. All transformants were analyzed by diagnostic PCR with location-specific primers and qRT-PCR (S1 Fig).

To induce *laeA* expression, the PCR fusion fragments of the *gpdA* promoter to the *laeA* ORF were digested with Hind III and KpnI and then cloned into the pPTRI vector (Takara, Japan) harboring the pyrithiamine (PT) resistance gene. The confirmed recombined vector was transformed into *Δspc105* and *A. flavus* NRRL3357 protoplasts. For *laeA* gene deletion, the 5′ and 3′ flank regions of *laeA* were amplified and fused with the *ptrA* gene, which was amplified from the pPTRI vector. The deletion construct was introduced into the *OE::spc105* and *A. flavus* NRRL3357 strains. Transformants were selected on glucose minimal medium (GMM) containing 0.1 mg/ml pyrithiamine (PT, Sigma) and were confirmed by PCR and qRT-PCR analysis (S1 Fig).

### Fungal physiology experiments

For morphological observation of colonies, strain spores were point inoculated on PDA, GMM, and Czapek’ medium (CZ, Difco) solid plates. For quantitative analysis of conidial production, PDA plates were overlaid with 5 ml of a suspension of conidia (10^6^ spores/ml) in 0.7% molten agar. Conidia were harvested from three 7-mm cores which were individually homogenized in 0.05% Triton X-100 solution and quantified with a hemocytometer. Sclerotia production was measured as previously described [62] by counting sclerotia from CZ culture plates after incubation for 14 days at 30°C in the dark.

### Microscopic analysis and nuclear staining

For the conidia germination assay, *A. flavus* conidia were inoculated in 5 ml potato dextrose broth (PDB, Difco) liquid media with coverslips at 37°C and 25°C. The morphology of germinated conidia and hyphae were observed using a light microscope at different time intervals. For examination of nuclear division, samples were fixed with an appropriate fixing solution for 10 min [63]. They were washed twice in distilled water and stained in 100 ng/mL 4′,6-diamidino-2-phenylindole (DAPI, Sigma). Samples were then washed twice in phosphate-buffered saline (PBS) and viewed using a Leica fluorescence microscope.

### Examination of AF, kojic acid, and CPA production

AFB_1_ production in mycelia and cultures were measured using modified thin layer chromatography (TLC) analysis as previously described [64]. The extract residue was resuspended in acetone and developed on Si250 silica gel plates with chloroform-acetone (9:1, v/v). The plates were visualized under 254-nm UV light. Standard AFB_1_ was purchased from Sigma.

Kojic acid production was determined using the colorimetric method [65]. Briefly, *A. flavus* strains were cultured on PDA supplemented with 1 mM FeCl_3_. Kojic acid forms a chelated compound with ferric ions and subsequently generates a red color, allowing for a qualitative comparison between different strains.

CPA production was detected using TLC [66]. The 50-mL PDB medium cultures were grown statically for 7 days and then harvested and extracted with chloroform. The TLC plate was sprayed with a 2% solution of oxalic acid in methanol followed by heating in a dry oven at 85°C for 15 min. Toluene–ethyl acetate–acetic acid (8:1:1, v/v/v) was used as the developing solvent. All the above experiments were performed with four replicates.

HPLC analysis of the above three secondary metabolites was performed using a Shimadzu LC-20AT system (Shimadzu). Extracts of each sample were separated on a Luna C18 column (Phenomenex) which was equilibrated with a running solvent consisting of acetonitrile–water (36:65) for AF detection, methanol–water (10:9) for kojic acid detection, and acetonitrile–0.1% trifluoroacetic acid (50:50) for CPA production.

### Peanut infection assay

Peanut cotyledon colonization assay was performed as described previously [17]. Surface sterilized peanut cotyledons were inoculated with a 10^5^ spores/mL of *A. flavus* and incubated for 3 days at 30°C in dark conditions. The infected seeds were collected in 50 mL Falcon tubes and mixed with 10 mL sterile 0.05% Triton X -100, followed by 1 min vortex to release the spores. The spores were diluted and counted using a hemocytometer. AFB_1_ production detection was performed by adding 10 mL chloroform to the Falcon tubes, followed by shaking for 5 min three times.

### qRT-PCR analysis

qRT-PCR assay was performed using the Applied Biosystems Step One Plus system (Invitrogen) with SYBR Green detection as described previously [62]. Gene expression levels were normalized (ΔΔCt analysis) to *A. flavus* β-actin gene expression levels. All analyses were performed in triplicate.

### RNA sequencing and data analysis

RNA samples from three *A. flavus* independent biological repeats were prepared. Strains were grown in PDB at 25°C for 48 h and mycelia were harvested immediately for RNA extraction using Trizol Reagent (Invitrogen). RNA quality and quantity were determined using an Agilent 2100 bioanalyzer system and RNA integrity number was calculated. Sequencing libraries were prepared with high-quality RNA samples (RNA integrity number ≥ 8) using an Illumina TruSeq RNA Sequencing Kit and sequenced on an Illumina Hiseq2500 system (Oebiotech, Shanghai, China).

Raw sequencing reads obtained for each sample were quality controlled using FastQC [67] and then filtered to remove the low-quality reads using version 2.3.3 of the NGS QC TOOLKIT [68]. The remaining reads were mapped to the *A. flavus* NRRL3357 genome (GCF_000006275.2) using version 2.0.13 of tophat2 [69] and version 2.2.4 of bowtie2 [70]. Gene expression level was normalized by calculating the number of Fragments Per Kilobase per Million reads mapped (FPKM) [71], and the base mean of each gene from the six sequenced samples was analyzed using DESeq software [72] to identify DEGs. Genes with |log2 (fold change)| ≥1.5 and adjusted *P* value (padj) ≤ 0.01 were defined as significantly differentially expressed. The gene-function annotation was conducted based on the GO and KEGG databases [73].

### Yeast two-hybrid assay

Y2H assay was performed using the Matchmaker Gold Y2H System (Clontech), according to the manufacturer’s instructions. The *spc105* and *laeA* inserts were amplified from *A. flavus* cDNA templates and cloned into pGADT7 and pGBKT7, respectively. The sequences of all inserts were verified by DNA sequencing analysis. Recombined vectors were transformed into the *S. cerevisiae* Y2HGold strain and plated onto selective media (SD/-Leu/-Trp). Then, the transformants were plated on the selective synthetic dextrose medium (SD/-Leu/-Trp/-His/-Ade/+X-α-gal) and incubated at 30°C for 3–5 days.

### Recombinant protein purification and GST pull-down assay

Heterologous expression and subsequent purification of Spc105 and LaeA proteins were conducted in *E. coli* BL21 (λDE3) using a combination of GST fusion vector pGEX4T-1 (GE Healthcare) and N-terminal 6X histidine-tag fusion vector pET28a. Briefly, the *spc105* gene was inserted into pGEX4T-1 to express GST-Spc105, which was subsequently purified on glutathione-agarose 4B (GE Healthcare) following manufacturer’s recommendations. The *laeA* gene was cloned into pET28a to yield a His_6_-LaeA fusion protein, which was purified using Ni-NTA agarose (GE Healthcare).

GST pull-down experiments were performed according to the manufacturer’s recommendations. Briefly, 20 μg purified GST-Spc105 protein, 25 μL glutathione magnetic beads (Pierce), and 20 μg purified His6-LaeA protein were co-incubated for 3 h at 4°C in PBS buffer. The magnetic beads were subsequently washed 6 times with PBS buffer and boiled for 10 min in SDS-PAGE loading sample buffer. SDS-PAGE and Western blot analysis were subsequently conducted. Immunodetection of His_6_-LaeA was performed using a Mouse His-tag monoclonal antibody at a dilution of 1:5000 (Proteintech) followed by Goat anti-Mouse HRP 1:10000 (Proteintech).

### Statistical analysis

All statistical analyses were performed using GraphPad Prism (version 5.0; GraphPad Software) and *P* ≤ 0.05 was considered a significant difference.

## Data availability

RNA-seq data were deposited in the NCBI Sequence Read Archive under accession number PRJNA498762.

## Acknowledgments

We are thankful to Prof. Yongjun Lu (School of Life Sciences, Sun Yat-sen University) for providing strains and plasmids used for Y2H and GST pull-down assays. We also thank Prof. Shihuang Wang (School of Life Sciences, Fujian Agriculture and Forestry University) for providing pPTRI vector.

## Supporting Information

**S1 FIG. Preparation of mutant strains used in this study.** (A) Diagram showed the replacement of *spc105* with the *pyrG* marker by a homologous recombination event. (B) PCR, RT-PCR and RT-qPCR verification of *Δspc105* and *OE::spc105* transformants. PCR verification of the four putative *spc105* deletion and two putative *OE::spc105* transformants was performed with primer pairs spc/NF and spc/NR and OE5F and OE3R; RT-PCR analysis of *spc105* expression in *Δspc105* and *OE::spc105* strains from 48 h cultures was performed using primer pairs RT-spcF and RT-spcR; Relative gene expression levels of spc105 in transformants were analyzed using ΔΔCT analysis. (C) Verification of eGFP-spc transformants. Left: Results of PCR amplification from control strain (NRRL3357-5) and two putative eGFP-spc transformants using primer pairs OE5F and OE3R; Middle: RT-PCR results of *gfp* gene expression in two putative transformants; Right: Western blot detection of GFP expression in eGFP-spc-14. (D) Verification of spc/laeA double mutants.

**S2 FIG. Hyphae dry weight measurement of spc105 mutant strains in PDB culture.** Strain spores were inoculated in 30 mL PDB broth and incubated with shaking at 200 rpm. Mycelia were harvested at indicated time points by vacuum filtration.

**S3 FIG. Overlaid chromatograms analysis of several secondary metabolite production in *A. flavus spc105* mutant strains.** (A) (B) and (C) represents AFB_1_ kojic acid, and CPA, respectively. Results confirmed that little to no AFB_1_ and extreme low level AFB_1_ is present in extracts of the *Δspc105* strain at 25°C and 30°C, respectively.

**S4 FIG. Global gene expression analysis of two samples with three replications.** (A) Cluster analysis of sample to sample to verify the similarities between parallel samples. (B) The boxplot of overall gene expression level of two samples with three replicates. (C) A volcano plot shows the total differentially expressed genes (DEGs) (red spots, P ≤ 0.01) between *Δspc105* and WT samples. (D) The heatmap showed the DEGs between *Δspc105* and WT samples. The data used to construct the heatmap was based on the log2 value of the FPKM values of all DEGs.

**S5 FIG. KEGG analysis of DEGs between *Δspc105* and WT.**

**S1 Table. Differentially expressed backbone genes in *Δspc105* relative to WT.** Genes with |log2 (fold change)| ≥1.5 and adjusted *P* value (padj) ≤ 0.01 were defined as significantly differentially expressed.

**S2 Table. Transcript levels relative to WT for AF cluster genes in *Δspc105*.**

**S3 Table. Differentially expressed oxidative stress response genes in *Δspc105* relative to WT.**

**S4 Table. *Aspergillus flavus* strains used in this research and their genotypes**

**S5 Table. Primers used in this study.**

